# Metabolism and chemical diversity evolve in response to pollinator availability

**DOI:** 10.64898/2026.02.13.702789

**Authors:** Elisabeth Authier, Léa Frachon, Jeanne Friedrichs, Lukas Brokate, Robert R. Junker, Caroline Müller, Thomas Dussarrat

## Abstract

Phytochemistry is a core player in shaping plant-pollinator networks and pollination services. Yet, little is known about the dynamic evolution of phytochemical traits in response to limited pollinator access, especially concerning chemical diversity. We combined an evolutionary experiment manipulating pollinator access with predictive metabolomics to uncover evolutionary changes in phytochemical traits of *Brassica rapa*. Our results unveiled chemical changes in both leaf and flower chemistry. Moreover, plants under selection by limited pollinator access showed a decreased chemical richness and diversity and a modulated primary and specialised metabolism, which could be used to predict pollinator access with 88% accuracy. Chemical indices and metabolites responding to pollinator access were associated with variation in flowering time and performance of outcrossing flowers. Our findings provide key insights into the influence of pollinator access on plant chemistry and indicate a risk of pollinator decline and losses of chemical diversity for plant-pollinator network structure and ecosystem dynamics.

## 1. Introduction

Plant-pollinator mutualistic interactions are fundamental to ecological networks (Bascompte & Jordano 2007), driven by a complex interplay of traits that have fostered remarkable floral diversity. Pollinators act as key selective forces impacting multiple plant traits to enhance reproductive success of flowering plants, from floral morphological and mating system-related traits (e.g. flower shape, reproductive organs, selfing ability) to chemical (e.g. volatile compounds, pigments, nectar composition, chemical defence) and ecological traits (e.g. phenological synchronisation, niche partitioning) (Parachnowitsch & Junker 2021). However, human activities threaten pollinators (Hallmann *et al*. 2017; Potts *et al*. 2010), resulting in profound changes in plant-pollinator interactions (Biesmeijer *et al*. 2006; Eckert *et al*. 2010; Hegland *et al*. 2009; Petanidou *et al*. 2014), and ultimately, in biodiversity and human food security (Aizen *et al*. 2009; Klein *et al*. 2007). Therefore, understanding the rapid adaptive potential of flowering plants in response to pollinator community composition shifts is crucial for predicting plant species persistence, effective wild plant species management, and the optimisation of plant breeding programs.

Although challenging, studying short-term adaptive dynamics offers unique opportunities to uncover the molecular bases of plant-pollinator interactions. For instance, to demonstrate phenotypic evolution in traits related to plant attractiveness under pollinator decline conditions, Thomann et al. (2015) regrew *Adonis annua* seeds from plants 18 years apart in a resurrection experiment. However, limited ancestral seed availability makes this approach incompatible with typical project timelines. Recent studies have performed experimental evolution experiments using fast cycling plants to uncover their adaptive potential to pollinator changes in controlled conditions (Gervasi & Schiestl 2017; Ramos & Schiestl 2019). Authier et al. (Authier *et al*. 2026) grounded this approach in ecological realism by using a common garden setting with natural pollinator communities. Their findings reveal rapid phenotypic trait changes in response to limited temporal access to pollinators. Resurrecting first and last generations helps to distinguish adaptive response to known selective agents from plastic effects, particularly in environmentally sensitive traits such as chemical signals involved in plant-pollinator interactions.

Phytochemicals act as a flagship component in the interactions of plants with their abiotic and biotic environment (Dussarrat *et al*. 2024; Whitehead *et al*. 2021). Variation in phytochemical traits can be studied at the scale of specific metabolites or chemical indices, which include richness, evenness and disparity of the different chemical classes (*e.g.* phenolics) (Petrén *et al*. 2024). At the metabolite level, floral scent is known to play a core role in plant-pollinator interactions (Burkle & Runyon 2019; Dudareva *et al*. 2006; Larue *et al*. 2016; Schiestl 2010). In addition, sugars, amino acids and specialised floral metabolites, which define flower fragrance and colour, can also influence pollinator preference (Borghi & Fernie 2017; Stevenson *et al*. 2017). The question of whether the adaptive response of plant individuals to variations in pollinator communities relates exclusively to their flowers or extends to leaf chemistry remains, however, poorly explored. Chemodiversity (*i.e.* chemical diversity) indices, such as richness and Shannon diversity of terpenes or glycoalkaloids influence herbivore behaviour and performance (Calf *et al*. 2019; Ziaja & Müller 2023), but also flower visitors (Hetherington-Rauth & Ramírez 2016; Sasidharan *et al*. 2024). However, these studies were limited to the analysis of specific chemical classes and well-known compounds, lacking a more holistic approach using untargeted metabolomics and including disparity, the third level of chemodiversity (Dussarrat *et al*. 2022; Petrén *et al*. 2023). Besides, while the mechanisms underlying the evolution and maintenance of chemical diversity are debated (Thon *et al*. 2024; Whitehead *et al*. 2021), analyses of the evolutionary response of plant metabolism to interaction partners remain scarce, particularly under the pressure of pollinator decline. Hence, combining an evolutionary experiment and resurrection approach with metabolomics has the potential to provide a unique understanding of the evolutionary chemical consequences of a decline in pollinator access.

Here, we explored the dynamic evolution of phytochemical traits in response to contrasting levels of pollinator temporal access in a realistic ecological context using the Brassicaceae *Brassica rapa* as a model system due to its short life cycle (Gervasi & Schiestl 2017). More specifically, we asked (i) which phytochemical traits are selected in flowers and leaves of *B. rapa* to cope with limited access to pollinators and (ii) whether the phytochemical traits (*i.e.* compounds or diversity indices of chemical classes) involved are linked to relevant morpho-anatomical traits such as the number of seeds per fruit. We hypothesised chemical changes in both organs, as previous studies reported the importance of functional traits that involve a general plant response (*e.g.* flower height) (Fornoff *et al*. 2017). When facing limited pollinator access, we also hypothesised a decrease in chemical diversity, as flowers might become more specialised to attract certain pollinators and therefore select specific compounds. Alternatively, plants could reduce their dependency on pollinators by decreasing their attraction and favouring selfing (Thomann *et al*. 2013). To test these hypotheses, six generations of fast cycling *B. rapa* were grown under contrasting conditions of pollinator access to flowering plants. In detail, we explored the variation in phytochemical traits between individuals from the first and sixth generations and tested the independence of leaf and flower chemical evolution. Furthermore, predictive metabolomics was employed to unveil a set of phytochemical traits that could predict pollinator access. Significant chemical indices and compounds were then linked to variation in phenotypic traits.

## 2. Materials and Methods

### 2.1. Design of the evolutionary experiment

To unravel the chemical mechanisms involved in the rapid evolutionary response of plants to limited temporal access to pollinators, we combined an experimental evolution experiment using *B. rapa* fast cycling (Wisconsin Fast Plants®) (Wendell & Pickard 2007) with a resurrection approach detailed in Authier et al. (Authier *et al*. 2026). Briefly, we carried out an experimental evolution in a common garden at the Botanical Garden of the University of Zurich (Switzerland) over six generations using three contrasting pollination treatments (see below). The experiment was conducted for two consecutive years (2021 and 2022) during the pollinator activity period (April to November). The starting population consisted of 108 full-sib seed families (hereafter called families), which we divided into three replicates of 36 families each. For each replicate, we applied all three pollination treatments (Fig. 1). After germination in greenhouse, individuals from each replicate and each treatment (hereafter called replicate population) were placed into nine outdoor cages in a common garden. Experimental evolution was repeated over six generations, with selection mediated by three distinct pollination treatments. The first treatment acts as the control treatment (latest resulting generation named G6 control) in which hand-pollination was performed in constantly closed cages (Fig. 1). We ensured that all plants were cross-pollinated by another plant within a given replicate. The second treatment (named G6 full) involved full plant access to the natural pollinator community at the Botanical Garden. Open flowers were marked using coloured paper tape, and the cages were opened on a single day during peak pollinator activity (from 11 am to 4 pm), ensuring that the entire pollinator community could visit plants. A single replicate cage was opened each day to avoid cross-pollination among replicates. Finally, in the third treatment (named G6 limited), the temporal access to the pollinator community was limited to one hour (Fig. 1). Each replicate cage was opened for one hour during the period of peak activity of pollinators (12 pm to 1 pm). As for the full access treatment, opened flowers were marked. We opened only one cage per day, and each cage was opened for just one day per generation. These treatments were repeated in each generation using the same method. Selection was applied using a fitness proxy for each generation, namely the product of the total number of seeds produced times the germination rate (detailed in(Authier *et al*. 2026)). To observe the evolutionary process and minimise environmental effects, which are highly sensitive factors in metabolic analysis, we resurrected seeds from the first and last generations in greenhouse. To reduce maternal effects, we regrew plants twice in the greenhouse and generated the second resurrected generation using hand-pollination. The second resurrected generation was sown in August, 3rd 2023 in Einheitserde CLASSIC® substrate (PATZER ERDEN GmbH, Sinntal-Altengronau, Germany) and plants placed into a phytotron for two weeks (constant temperature 21°C day and night, 16h light cycle, and 60% of relative humidity). One week later (August 11th 2023), plants were potted into individual pots (7×7 cm) in standard soil and moved to the greenhouse (constant temperature 21°C, 16h light and 60% relative humidity) of the Irchel Campus (University of Zürich). The phenotypic traits measurement, as well as tissue sampling for metabolomic analysis, were performed on this second resurrected generation (for more details see (Authier *et al*. 2026)).

**Fig. 1.**
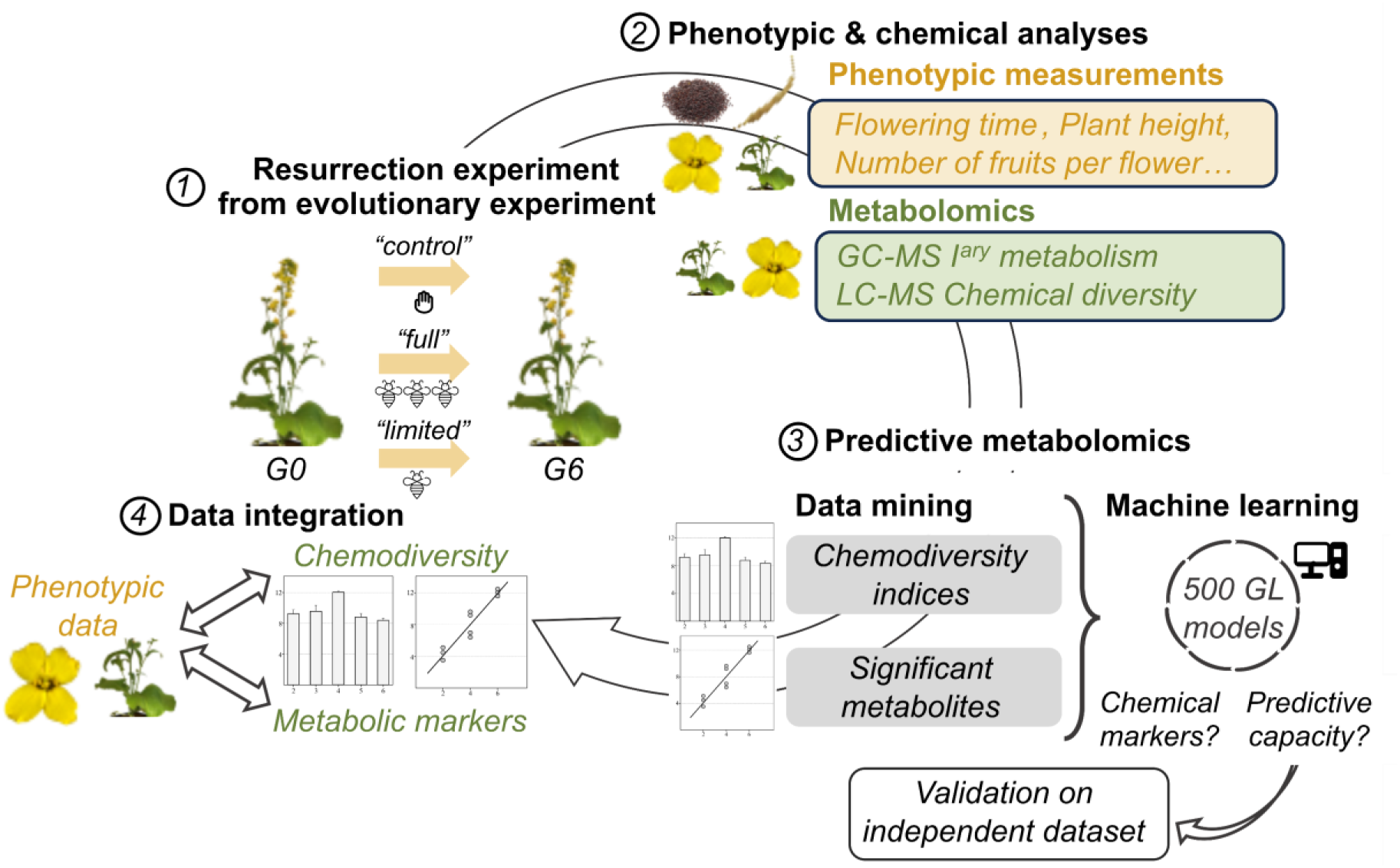
| A simplified scheme of the experiment. Six generations of *Brassica rapa* were developed under three distinct treatments as follows: hand pollination and full or limited access to pollinators (indicated with a hand and 3 or 1 pollinators in the graph, respectively). Leaf and flower samples were collected on plants from generations 0 and 6 after the resurrection experiment. *G0 and G6*: generation 0 and 6. Images pollinators and hands represent the three treatments (*i.e.* hand pollination, full access or limited access to pollinators). *GC/LC-MS*: gas or liquid chromatography mass spectrometry. *GL models*: generalised linear models.

### 2.2. Phenotypic data

Phenotypic traits related to phenology (flowering time), plant architecture (number of branches and plant height), and fitness traits were measured in the resurrection experiment across the first and last generations. The ‘flowering time’ was defined as the number of days between sowing and the appearance of the first flower. The ‘number of branches’ and ‘plant height’ were counted and measured at the end of the plant life cycle. Finally, traits related to fitness and plant mating system were assessed. To assess outcrossing ability, three flowers were hand-pollinated using pollen mixture from all individuals within the same replicate population. Additionally, we hand-pollinated three flowers with pollen from the same individual to evaluate selfing ability (autogamy). At plant maturity, the number of fruits produced by the three hand cross-pollinated flowers (referred as ‘Number of fruit outcrossing’), and the three hand self-pollinated flowers (referred as ‘Number of fruit selfing’) were counted. For fruits from hand cross-pollinated flowers, the fruit length was measured and the mean among these three fruits was calculated (referred as ‘Fruit length outcrossing’). The ‘Fruit length selfing’ (from self-pollinated flowers) was defined similarly. Finally, the mean of seeds per fruit was calculated for both hand cross-pollinated flowers (referred as ‘Number seed/fruit outcrossing’) and hand self-pollinated flowers (referred as ‘Number seed/fruit selfing’).

### 2.3. Sampling

From the initial stock of 36 “G0”, X “G6 control”, 36 “G6 full”, and 36 “G6 limited” individuals, 20 individuals per replicate population were randomly selected for sampling (Authier *et al*. 2026). From these populations, samples were collected only if the number of flowers was sufficient for the measurements of phenotypic traits. Leaves and flowers were collected during flowering time on August 29^th^ 2023 between 1:15 pm and 4:15 pm. Each leaf and flower replicate consisted of the two youngest leaves and six flowers per individual, respectively. Samples were collected, snap-frozen into liquid nitrogen and stored at –80°C until freeze-drying (Alpha 2-4 LSC (Martin Christ, Osterode am Harz, Germany) coupled to a Pfeiffer DUO 10 M vacuum pump (Pfeiffer Vacuum, Aßlar, Germany) for two days). For some samples, the quantity of leaves or flowers was not sufficient for chemical analyses, in which case powders from a same replicate and treatment were taken (e.g. sample 197B included powder from two plants from generation 6 under limited pollinator access, see Table S1). A total of 195 leaf and 182 flower samples were used for chemical analyses, which included 50 leaf and 44 flower samples for G0, 52 leaf and 47 flower samples for G6 control, 49 leaf and 49 flower samples for G6 limited, 44 leaf and 42 flower samples for G6 full (Table S1).

### 2.4. Metabolite extraction

For the analysis of primary compounds, 8 mg of dried flower or leaf material were extracted in a mixture of 1:2.5:1 (v:v:v) chloroform, methanol, Millipore water with ribitol (99%, AppliChem) as an internal standard as previously described (Schweiger *et al*. 2023). Derivatisation and silylation were performed using methoximation with *O*-methylhydroxylamine hydrochloride (> 97%, Sigma Aldrich-Merck; >98%, Alfa Aesar) in pyridine (≥ 99.9%, HPLC grade; Honeywell) and *N*-methyl-*N-*trimethylsilyl-trifluoracetamide (> 98%; Macherey-Nagel).

For metabolic fingerprinting, a 90% methanol solution (v:v) with hydrocortisone (10 mg/L) as internal standard (Sigma-Aldrich, Steinheim, Germany) was added to 10 mg of dried homogenised powder and sonicated in an ice bath for 15 min. After centrifugation, supernatants were collected and filtered using 0.2 µm syringe filters (Phenomex, Torrance, CA, USA).

### 2.5. Metabolomics

Primary compounds were analysed via GC-MS (GC-2010 Plus coupled to QP2020; Shimadzu). A VF-5 ms column (30 m x 0.25 mm x 0.25 µm, Agilent Technologies) was used with an injection temperature of 225°C, a helium flow of 1.14 mL/min and a split of 10. The temperature gradient included an oven temperature of 80°C for 3 min followed by a first ramp of 5°C/min to 310°C held for 2 min, a second ramp of 15°C/min to 325°C held for 3 min, and an ion source at 230°C, as previously described (Schweiger *et al*. 2023). Measurements were performed in electron impact positive ionisation mode at 70 eV. Retention indices were determined using an alkane standard mix (C7-C40, Sigma Aldrich). Data analysis was performed on GC-MS Postrun Analysis (v. 4.45, Shimadzu). Retention indices and mass spectra were screened on chemical databases (in-house database of primary metabolites, GOLM Metabolome database, National Institute of Standards and Technology) (Kopka *et al*. 2005; Lemmon *et al*. 2010; Schauer *et al*. 2005) for compound annotation, as previously described (Schweiger *et al*. 2023). Peak areas were divided by the area of the internal standard for each sample, and the mean blank area was subtracted from each compound. Finally, peaks were related to sample dry mass and compounds with two analyte peaks were summed, yielding 37 detected and annotated compounds (Table S4).

Metabolic fingerprints were analysed using UHPLC-QTOF-MS/MS (UHPLC: Dionex UltiMate 3000, Thermo Fisher Scientific, San José, CA, USA. QTOF: compact, Bruker Daltonics, Bremen, Germany). A Kinetex XB-C18 column (150 x 2.1 mm, 1.7 µm, with guard column; Phenomenex) was used at 45°C with a 2 to 30% gradient from Millipore-H2O-0.1% formic acid (FA) to acetonitrile-0.1% FA within 20 min and increase to 75% acetonitrile-0.1% FA within 9 min (flow rate of 0.5 mL/min). Negative electrospray ionisation was used at a spectra rate of 6 Hz (*m/z* range of 50-1300) with optimised MS parameters, including capillary voltage 3000 V, end plate offset 500 V, quadrupole ion energy 4 eV and collision energy 7 eV (Dussarrat *et al*. 2023). MS/MS spectra were obtained using the AutoMS/MS mode. The same protocol was used for leaf and flower samples. Pre-processing was performed using the T-ReX 3D algorithm of MetaboScape (v. 2021b, Bruker Daltonics) with the following settings: features in minimum 3 samples, intensity threshold 1000, minimum peak length 11, MS/MS import method Maxsum. Subsequent data cleaning (average quality control (QC) intensity 5 times higher than average extraction blank intensity (i.e. peak height), coefficient of variation in QC < 30%) yielded 846 and 1780 ions for leaves and flowers, respectively. The relatively low quantity of ions detected in leaves is explained by the fact that the spectra rate had to be adjusted to avoid overloading peaks in flowers. LC-MS datasets were normalised using median normalisation, cube-root transformation and Pareto scaling on MetaboAnalyst v.5 (Pang *et al*. 2021) prior to statistical analysis. Non-normalised LC-MS datasets are available in Tables S5 and S6.

Chemical class annotation was performed on SIRIUS (v. 6.1.0) with Q-TOF default parameters (10 ppm and de novo plus bottom up molecular formula generation with a threshold *m/z* of 500). Metabolite structure and chemical class predictions were performed with CSI:FingerID and CANOPUS with a confidence score defined using “approximate” mode and the Natural Products Classifier (NPC) ontology (Kim *et al*. 2021). PubChem was not used as a reference. For each feature, the most probable NPC pathway, superclass and class were assigned only if the confidence score was ≥ 0.8, as recommended (Hoffmann *et al*. 2022) (Table S7). Next, richness and Shannon diversity indices were defined for each chemical pathway, superclass and class using the R package *chemodiv* (Petrén *et al*. 2023). To consider disparity, the cosine similarity score between MS/MS spectra was assessed via the Global Natural Product Social Molecular Networking (GNPS, https://gnps.ucsd.edu/ProteoSAFe/static/gnps-splash.jsp) platform and used to define the dissimilarity matrix (1 – spectral similarity, *i.e.* 1 – cosine score) and therefore calculate the functional Hill diversity for each chemical class. In total, 85 and 98 indices were obtained for leaves and flowers, respectively. Prior to normalisation (median normalisation, log transformation and autoscaling) and statistical analysis, superclasses and classes with less than 4 compounds (0.5% of leaf processed ions) were removed from the dataset, as the biological meaning of their analysis might be questionable (Brückner & Heethoff 2017; Petrén *et al*. 2023). However, it is interesting to note that no precise threshold has yet been defined in the literature, and all detected classes are usually considered (Defossez *et al*. 2021). The non-normalised “chemical indices” datasets for leaves and flowers are available in Tables S8 and S9.

All non-normalised and normalised datasets (GC-MS, LC-MS and chemical indices) are available online (see Data availability section).

### 2.6. Statistical analyses

As a first step to explore the phytochemical traits modulated in flowers and leaves of *B. rapa* in response to limited access to pollinators, we tested the correlation between leaf and flower LC-MS datasets using the Mantel test (*vegan* package with “Kulczynski” method) and two-way orthogonal partial least square analysis (O2PLS, *OmicsPLS* package) in R (v. 4.2.1) (Bouhaddani *et al*. 2018; Oksanen *et al*. 2022; R Core Team 2022) (Table S10). The *vegan* package was also used to calculate Bray-Curtis dissimilarities between replicates and treatments with the “bray” method.

Subsequent statistics aimed to test the variation in phytochemical traits between sample groups. Each of the “LC-MS” and “chemical indices” datasets was divided into subdatasets 1 (*i.e.* subdataset 1 “LC-MS”, subdataset 1 “chemical indices”), which included samples from replicates 1 and 3, and subdatasets 2, which included samples from replicate 2 (Table S3). While variable selections were performed on subdatasets 1, subdatasets 2 were used for validation. The response of phytochemical traits to variation in access to pollinators was evaluated via analysis of variance (ANOVA) in MetaboAnalyst 5.0 (*P* < 0.05, FDR) (Pang *et al*. 2021). Principal component analyses (PCA) and partial least squares discriminant analyses (PLS-DA) were also performed in MetaboAnalyst 5.0, while heatmaps were developed using the package *pheatmap* (Pearson correlation, Ward algorithm) in R (v. 4.2.1) (Kolde 2019).

To test the capacity of either chemical indices or LC-MS features to predict the degree of pollinator access, generalised linear models (GLM) were conducted in R (v. 4.2.1) using the *glmnet* package (Friedman *et al*. 2010; Tay *et al*. 2023), as previously described (Díaz *et al*. 2024; Dussarrat *et al*. 2022) (Fig. 1). Briefly, subdataset 1 (for instance subdataset 1 LC-MS) was divided into a training (70%) and a validation (20%) set using stratified sampling to develop the model equation, which was then applied to the testing set (10%) to perform real predictions. To define the equation, binomial and multinomial models were developed using Ridge, Lasso and Elastic net regression to modulate the number of variables used in the models with a thousand penalty values ranging from 0 to 1, and cross-validation was applied to limit overfitting. Prediction accuracy was assessed on predictions performed on the testing set. Multinomial and binomial models were run 500 and 100 times respectively to cope with random partitioning. To statistically validate the models, the likelihood of spurious predictions was assessed using 100 or 500 permuted datasets, in which the sample classes (*e.g.* G6 limited) were randomly assigned to the samples. Model performances were compared using Tukey’s test (*P* < 0.01). For models based on LC-MS data, the top 5% metabolic predictors (89 ions for both leaves and flowers for comparability reasons) were extracted based on their occurrence in the models to assess their predictive capacity. Subsequently, the predictive capacity of the best phytochemical traits extracted from subdatasets 1 was tested on subdatasets 2 using the compounds that responded significantly in both datasets (117 and 141 for flowers and leaves, respectively) (Dussarrat *et al*. 2022). Barplots, boxplots and circos plots (Pearson correlation) were designed using *ggplot2, Hmisc* and *circlize* packages (Gu *et al*. 2014; Jr 2023; Wickham 2016).

## 3. Results

### 3.1. Diversity of multiple chemical families is influenced by pollinator access to plants

As a first step, we explored the variance in leaf and flower LC-MS datasets. Although leaf and flower LC-MS datasets were significantly correlated (Mantel’s test, P < 0.05), only 16.7% of the variance observed in the leaf dataset was explained by the flower dataset (and 5% in the reverse direction, Table S10). Besides, floral chemical indices (*e.g.* Shannon diversity of glucosinolates) only weakly reflect the chemical indices of the leaves. Yet, 14 correlations were observed between the two organs, including Shannon diversity of glucosinolates and shikimates and phenylpropanoid pathways, or total (*i.e.* all chemical classes) functional Hill diversity (Fig. S1). Next, a comparison of detected ions between treatments showed that *B. rapa* did not evolve new compounds in either leaves or flowers in response to limited pollinator access in six generations (Fig. S2). Unsupervised analyses could not efficiently discriminate individuals grown under full and limited pollinator access (Figs. 2 and S3). Although not significant, a distinct pattern was observed with PLS-DA in both organs where the “G6 limited” individuals differed slightly from the other treatments, suggesting the response of certain phytochemical traits (Fig. S3).

**Fig. 2.**
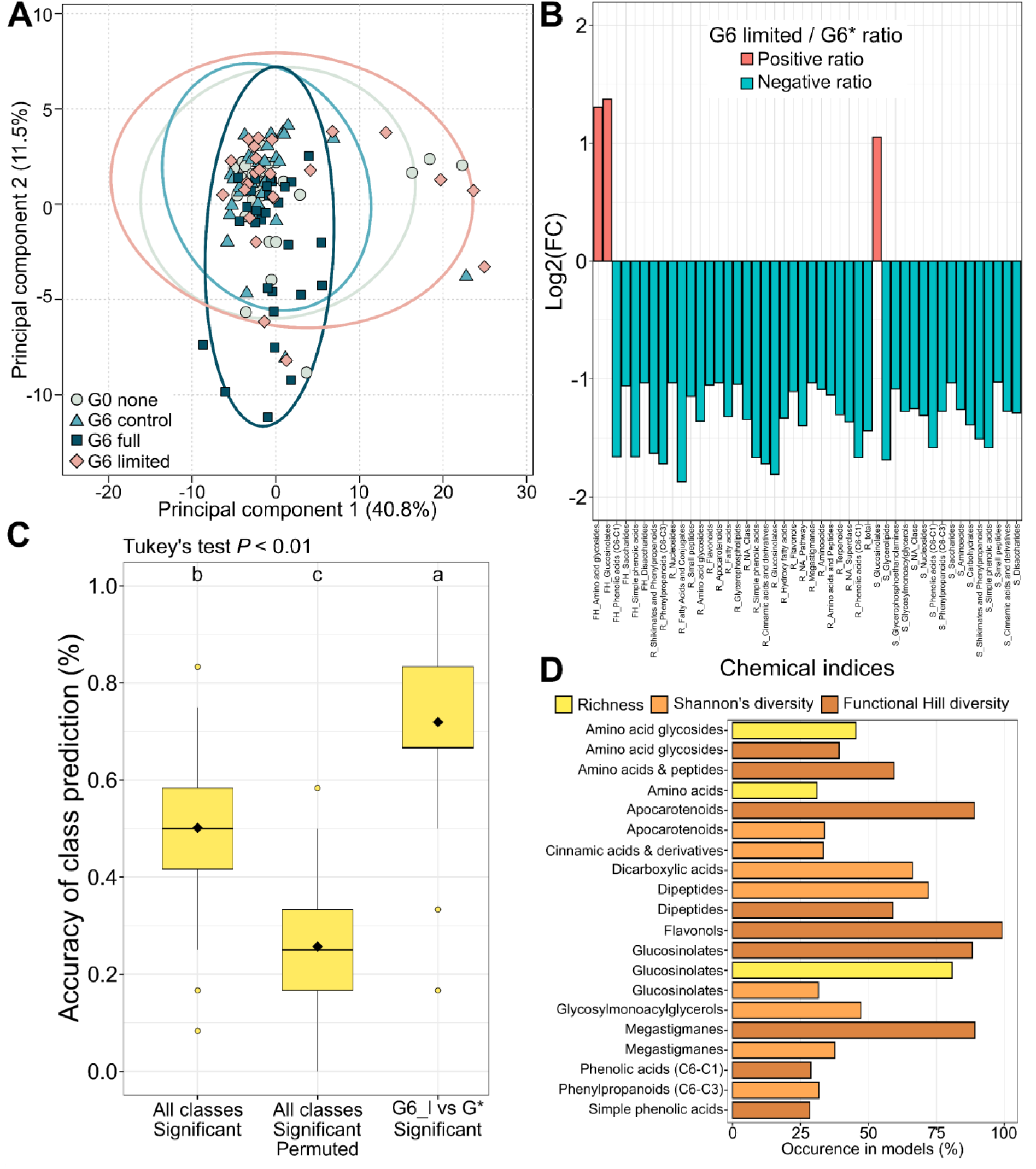
| Rapid evolution of chemodiversity in leaves and flowers of *B. rapa*. **A.** Principal component analysis on leaf samples using chemical indices. **B**. Fold change (G6 limited / G6*, which includes G6 control and G6 full) of significant indices in flowers (*P* < 0.5 FDR correction). Only indices with a fold change superior to 2 were represented. **C**. Box plot of the accuracy from 500 models evaluating the capacity of the 69 significant indices to predict classes “G6 limited” or “G*”, which includes G0 none, G6 control and G6 full. Letters indicate statistical difference between predictive capacity (Tukey’s test, *P* < 0.01). The square and the horizontal line show the mean and median, respectively. The box illustrates the lower, median and upper quartile values while circles show potential outliers. **D**. Ranking of the chemical indices in flowers based on their occurrence in generalised linear models. Only the top 20 predictive indices are illustrated in alphabetic order (see Table S8 for the exhaustive ranking list).

In dataset 1 (*i.e.* individuals from replicates 1 and 3), 69 floral chemical indices (*e.g.* richness flavonoids, total Functional Hill diversity) responded significantly (*P* < 0.05, FDR) to variation in pollinator access, compared with 3 indices for leaves (Table S11). Interestingly, richness and diversity (Shannon or Functional Hill diversity) indices decreased in individuals from G6 limited conditions for almost all chemical families except glucosinolates (the superclass “Amino acid glycosides” includes glucosinolates in NPC, Fig. 2). Next, we tested the capacity of chemical indices to predict pollinator access (Fig. 1). While the chemical indices were not able to discriminate very efficiently (50%) all groups (*e.g.* individuals in the G6 full or control groups), the binomial models achieved an accuracy of 72%, illustrating the ability to discriminate individuals with limited access to pollinators from other plants (Fig. 2). Models were statistically validated by the average accuracy of 26% obtained with 500 permuted data sets. A great proportion of the most predictive chemical indices referred to functional Hill diversity indices (9 of the top 20) and relevant chemical families included glucosinolates (amino acid glycosides and glucosinolates), phenolics (*e.g.* flavonols, phenolic acids) and terpenes (*e.g.* apocarotenoids) (Fig. 2 and Table S12).

### 3.2. Variation in primary metabolism

The response of 37 compounds from primary metabolism to variation in pollinator access was then tested using GC-MS. In dataset 1, nine and four compounds responded significantly (*P* < 0.05) in flowers and leaves, respectively (Table S13). After six generations under limited pollinator access, flowers showed lower concentrations of certain sugars such as sucrose and glucose and higher levels of other compounds such as threonate and 1,4-lactone-threonate (Fig. 3). Besides, some amino acids (*i.e.* glycine, threonine) and mannitol also responded significantly. Unfortunately, the number of samples present in datasets 2 did not allow biological validation of the importance of these primary compounds, as only three samples were available for the “G6 limited” group (Table S13).

**Fig. 3.**
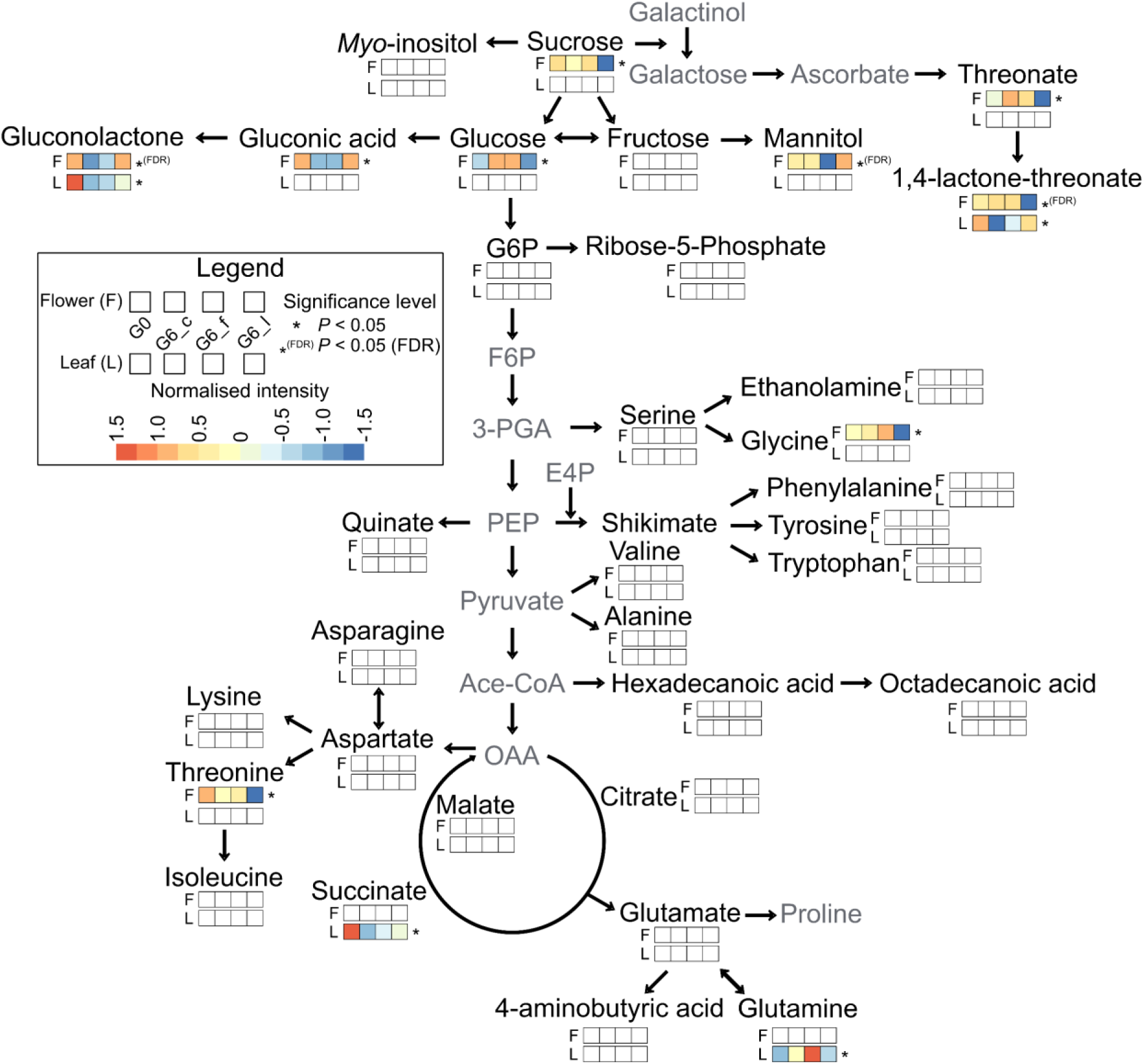
| Response of primary metabolism to the variation in pollinator access in leaves and flowers. Compounds in grey front were below the detection limit and were therefore not analysed in this study. This figure is a simplified representation of plant primary metabolism and does not include all reactions.

### 3.3. Evolution of *B. rapa* metabolism to limited pollinator access

Next, we explored the evolutionary response of metabolic fingerprints to limited pollinator access at the compound level. In dataset 1, 431 and 408 metabolic features responded significantly (*P* < 0.05, FDR) in flowers and leaves, respectively. Besides, while no variation in chemical indices was observed, the significant variation in 408 features showed that pollinator access also influenced leaf chemistry (Fig. 4A, 4B and Table S14). Evolutionary conditions were considerably predictable, with an average accuracy of 82% and 71% in flowers and leaves, respectively (Fig. 4C). Predictions were significantly different from the ones performed on permuted sets. Interestingly, the results of these multinomial models suggested a capacity to discriminate between G6 control and G6 full treatments, which was also supported by the heatmaps. Binomial models (*i.e.* comparing metabolic features of “G6 limited” individuals to individuals from other groups) reached 93% and 96% accuracy in flowers and leaves when using the best 5% predictors (89 features, Fig. 4C), validating the rapid evolution of *B. rapa* metabolism in response to limited pollinator access.

**Fig. 4.**
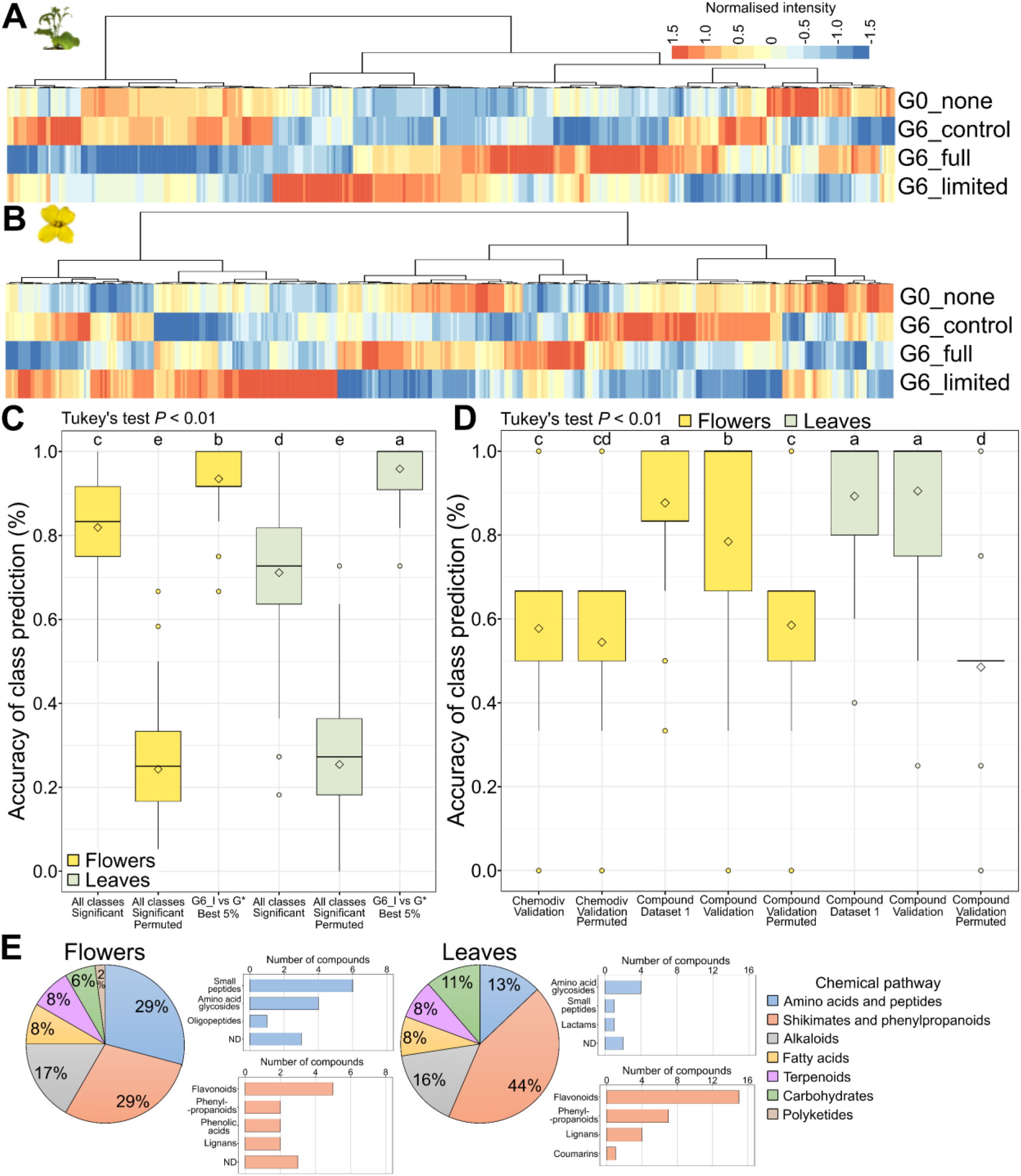
| Evolutionary response of leaf and flower metabolism to limited pollinator availability in *B. rapa*. **A-B.** Heatmaps (Pearson correlation, Ward algorithm) of the significant 408 and 431 significant features in leaves (**A**) or flowers (**B**), respectively (Tukey’s test, *P* < 0.05 FDR correction). **C**. Accuracy of the models. Multinomial models tested the capacity of flower and leaf metabolism to distinguish between “All classes” while binomial models tested pairwise comparisons between G6_limited (*G6_f*) and other classes (*G\**, which includes G0_none, G6_control and G6_full). Best 5% means that only the top 5% predictors (89 ions for both organs for comparability reasons) were used for the corresponding models. **D**. Accuracy of the validation models. For validation, predictive capacity of the 117 and 141 significant features (in both datasets) was tested on dataset 2 in flowers and leaves, respectively. Dataset 1 was reused to verify the predictive capacity of these selected features and permuted datasets were used to test the likelihood of spurious predictions. Significant variation in predictive score between modelling condition were accessed using Tukey’s test (*P* < 0.01). **E**. Chemical families represented among the 117 and 141 markers.

To provide an additional validation level, we verified the significance of the predictive chemical traits in the independent datasets 2, which is composed of plants from replicate 2 (Table S1). From the 69 significant floral chemical indices in dataset 1, six responded significantly in dataset 2 (*P* < 0.05) but none when considering FDR correction (Table S11). In leaves, while only three chemical indices were significant in dataset 1, 12 were significant in dataset 2 (*P* < 0.05, FDR). At the metabolite level, 37 and 18 were significant in both datasets in flowers and leaves, respectively (Table S14). Without FDR correction, 117 and 141 were significant in both flowers and leaves, respectively. The capacity of the significant chemical traits to predict pollinator access in dataset 2 was then tested (Fig. 4D). The predictive capacity of chemical indices in flowers could not be validated on dataset 2, as shown by the insignificant difference between the model accuracy of real and permuted datasets. This result may be explained by the dissimilarity of the samples from replicate 3 with the other replicates (Fig. S4). In contrast, average accuracy reached 88% and 78% in flowers for dataset 1 and 2, respectively and 89% and 91% for leaves, validating the predictive capacity of the significant features (Fig. 4D and Table S15). Notably, the capacity of leaf and flower metabolism to discriminate pollination type (*i.e.* by hand for G6 control or by pollinators for G6 full) was also validated in both datasets (Fig. S5). Interestingly, these predictive features mainly referred to flavonoids and amino acid glucosides, supporting their role in the response to a reduction in pollinator access (Fig. 4E and Table S16). Other chemical pathways represented by the predictive compounds included alkaloids, fatty acids, terpenoids and carbohydrates for both flowers and leaves. The main difference between the organs lies in a higher representation of amino acid derivatives and peptides, including glucosinolates, in flowers, compared with shikimates and phenylpropanoids, including flavonoids, in leaves.

### 3.4. The response of chemical markers is correlated with variation in phenotypic traits

Variation in pollinator access influenced various phenotypic traits, which included the number of fruits, the number of seeds per fruit, the number of branches and the flowering time (Fig. S6). For instance, plants under limited pollinator access showed a lower number of seeds per fruit from outcrossing flowers, as well as a lower fruit length, thus promoting a more autonomous reproductive strategy. The reduced chemical diversity (but increased for glucosinolate-related compounds) observed in flowers was mostly associated with an increase in flowering time under limited pollinator access. At the metabolite level, predictive leaf compounds were mostly correlated with the number of seeds per fruit (22 correlations with *P* < 0.05, FDR), the fruit length (33) and the number of branches (35) of *B. rapa* plants (Fig. 5). Besides, the number of floral predictive compounds correlated with flowering time (19 correlations with *P* < 0.05, FDR), number of seeds per fruit (22), number of fruits (17), fruit length (24) in flowers, and supported the relevance of these phenotypic traits for the response to variation in pollinator access (Fig. 5 and Table S17).

**Fig. 5.**
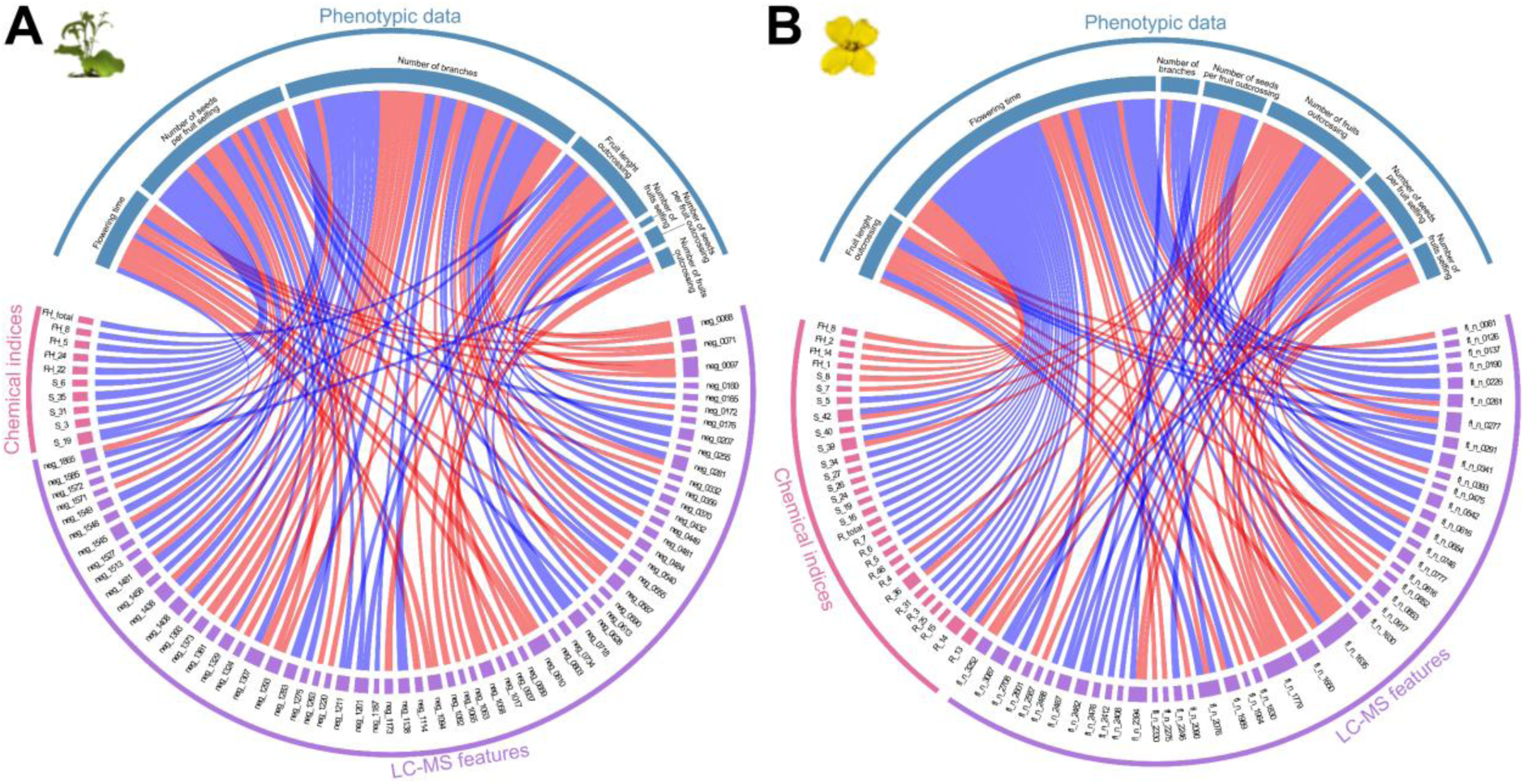
| Correlation between predictive chemical features and phenotypic variation. **A-B.** Circos plot representing the significant correlations (*P* < 0.05, FDR, Pearson correlation) between chemical indices or LC-MS features with phenotypic data in leaves (**A**) and flowers (**B**). Red links represent positive correlations, while blue links represent negative correlations. For better clarity, names of chemical families have been replaced with numbers, but the correspondence can be found in Table S14.

## 4. Discussion

Plants face a loss of pollinator diversity and abundance worldwide (Artamendi *et al*. 2024; Balfour *et al*. 2018). To adapt, flowering plant species may modulate flower functional traits such as flower reflectance and morphology, which influence the structure of plant-pollinator networks in natural ecosystems (Aguirre & Junker 2024; Fornoff *et al*. 2017). Phytochemical traits mediate variation in functional traits, and previous studies explored the influence of well-known floral metabolites on pollinator communities (Borghi & Fernie 2017; Hetherington-Rauth & Ramírez 2016; Stevenson *et al*. 2017). Yet, the capacity of chemical diversity to rapidly evolve to limited pollinator access has, to our knowledge, not been investigated, nor has the adaptive response of both flower and leaf metabolic fingerprints in a realistic ecological context (Parachnowitsch & Manson 2015). Here, we provided a holistic understanding of the evolutionary response of phytochemical traits to variation in pollinator access, highlighting the rapid evolution of major chemical classes that could be used to efficiently predict pollinator access in Brassicaceae.

### 4.1. The rapid evolution of *B. rapa* chemistry in response to limited pollinator access is not limited to flowers

Our results provide evidence that selective pressure on flowers not only shapes the evolution of flower chemistry but also that of vegetative organs, since flower and leaf chemistry could be used to predict pollinator access with 88 to 89% accuracy, respectively. This predictive accuracy supports the excellent integrative capacity of plant metabolism (Dussarrat *et al*. 2022) and is particularly interesting when considering that experimentally induced metabolic changes were generic to all studied *B. rapa* populations, suggesting stability across genetic backgrounds. Besides, the responsiveness of both flowers and leaves is consistent with the fact that, although studies have mainly focused on floral traits, vegetative parts have also exhibited a decisive influence on plant-pollinator interactions, such as plant height (Aguirre & Junker 2024; Zu & Schiestl 2017). In addition, flower and leaf chemistry under different regimes of pollinator access were correlated in our study, but only poorly co-varied between organs, highlighting the relevance of analysing the response of both organs. Similarly, with only 14 correlations, chemical indices of each chemical class in leaves poorly mirrored the ones in flowers (*e.g.* Shannon diversity of terpenoids in leaves was not linked to Shannon diversity of terpenoids in flowers), supporting the compartmentalisation of chemodiversity also previously observed between leaves and roots in *Tanacetum vulgare* (Kleine & Müller 2013). Finally, the response of certain phenotypic traits has shown that *B. rapa* plants favour selfing (*e.g.* decrease the number of fruit or the number of seeds per fruit from crossing flowers) in response to limited pollinator access, as shown previously (Authier *et al*. 2026; Ramos & Schiestl 2019), and the best leaf-predictive metabolites were associated with variation in these traits (in addition to the number of branches). Hence, this study underlines that the evolutionary response of plants to limited pollinator access is not limited to flower chemistry but rather involves a reorchestration of both leaf and flower metabolism. It remains to be tested which consequences such reorchestration of the leaf metabolism may have in an ecological context (*e.g.* influence on herbivores and (thus that turn into) florivores) (McCall *et al*. 2018; Rusman *et al*. 2022).

### 4.2. Flower chemodiversity is tailored to pollinator communities

Understanding how chemodiversity arises and evolves is essential, given its ecological and economic importance, but remain a major challenge (Thon *et al*. 2024). Here, we showed that limited pollinator access to plants leads to significant changes in chemical diversity indices of multiple chemical pathways. *B. rapa* flowers showed a decreased total chemical richness (*i.e.* all chemical classes) but also richness and diversity of multiple chemical classes such as phenolics and terpenoids under limited pollinator access, which supports our hypothesis. Pollinator communities are shaped by flower diversity in natural ecosystems, where higher floral chemodiversity leads to more generalised pollinator communities (Aguirre & Junker 2024; Fründ *et al*. 2010; Hanusch *et al*. 2025). The limited pollinator access imposed by our experimental design is associated with more specialised (*i.e.* less diverse) pollination (Authier *et al*. 2026), thus explaining the evolution towards a more specific – and thus reduced – chemodiversity presented to pollinators. This decreased chemodiversity is also consistent with the reduced outcrossing reproduction. Besides, a great proportion of the best predictive chemical indices, which allowed predicting pollinator access with 72% accuracy, referred to functional Hill diversity indices, supporting the role of chemical disparity, a dimension yet poorly explored in chemical ecology, in ecological processes (Petrén *et al*. 2023). This excellent predictive ability of chemical indices could not be observed on an independent dataset. This lack of consistency may be explained by the relatively low number of generations (six), which may not have been sufficient to consistently fix this evolutionary response in all the different *B. rapa* populations. Moreover, in our experimental evolution, significant changes in pollinator communities can be observed between spring, summer, and autumn on *B. rapa*, providing some natural variation in selection pressures by distinct pollinators between generations (Burkle & Runyon 2019). Although time-consuming, running a similar experimental design with one generation per year in midsummer may lead to a more pronounced and specific effect on phytochemical traits. This hypothesis is also supported by the fact that variation in chemical fingerprints could discriminate plants pollinated by hand (“G6 control”) from plants pollinated by natural pollinators (“G6 full”). Notably, the variation in chemical fingerprints between *B. rapa* plants of these treatments is also consistent with previous evolutionary experiments that have shown variation in phenotypic traits between hand-versus bee-pollinated plants in *B. rapa* (Dorey & Schiestl 2024).

### 4.3. Pollinator access shapes major chemical pathways from metabolite to chemical diversity

Our results indicated that pollinator access influenced the rapid evolution of primary and specialised metabolites, qualitatively and quantitatively. First, plants subjected to limited pollinator access displayed lower levels of some sugars (*e.g.* sucrose) and higher levels of other carbohydrates (*e.g.* gluconic acid, mannitol) in flowers than individuals from G6 control and G6 full treatments. These results exemplify the complex regulation of primary metabolism in flowers as multiple floral traits (*e.g.* pollen constitution, floral diameter) might respond to variation in pollinator access (Basnett *et al*. 2025; Parachnowitsch & Manson 2015; Thomann *et al*. 2013). Interestingly, limited pollinator access also led to lower levels of threonate and threonate-1,4-lactone, which are linked to ascorbate degradation (Truffault *et al*. 2017). This result supports the influence of redox state in pollination processes and may be the consequence of lower pollination in “G6 limited” *B. rapa* plants, as ascorbate levels were previously linked to pollen and flower development in *Solanum lycopersicum* (Deslous *et al*. 2021).

Predictions of pollination treatments using chemical indices highlighted the importance of various chemical pathways such as phenolics, terpenoids, and glucosinolates. The role of these classes was supported by their consistent occurrence among the best predictive metabolites. In leaves, a high occurrence was observed for the pathways “shikimates and phenylpropanoids” (44%), which includes flavonoids and phenolic acids, “amino acids and peptides” (13%), which includes glucosinolates, and alkaloids (16-17%). Additional classes were included in predictive chemical classes such as terpenoids and apocarotenoids in both organs. In plants, terpenoids are key players in plant-pollinator interactions (Sasidharan *et al*. 2024), while the observed importance of apocarotenoids functional Hill diversity in our study could be a result of their role as colour pigments or their link with phytohormones via carotenoid cleavage (Imtiaz *et al*. 2023). Besides, a higher occurrence (29% of most predictive compounds) of “amino acids and peptides” (and especially glucosinolates) and a lower occurrence of “shikimates and phenylpropanoids” (29%) were observed in flowers compared to leaves. In flowers, the role of glucosinolates in plant-pollinator interactions is widely recognised (Hopkins *et al*. 2009). The increase in glucosinolate diversity under limited pollinator access observed in our study is consistent with the decrease in the efficiency productivity of outcrossing flowers. In fact, plants that favour selfing will depend less on pollinator attraction and could therefore increase plant defence mechanisms against herbivores or florivores, for instance (Dorey & Schiestl 2024; Ramos & Schiestl 2019; Sasidharan *et al*. 2023). The rapid evolution of phenolics highlighted in flowers and leaves at both chemical diversity and metabolite levels supports their role in mediating plant-pollinator network structure in natural ecosystems. The response of phenolics is in line with previous observations that pollinators are the main driver for flower colour evolution (Tenhumberg *et al*. 2023). In fact, phenolics influence flower attractiveness by adjusting flower colours with carotenoids (Hoballah *et al*. 2007) and are also known to influence the evolution of pollination mutualism, especially when interacting with sugars in nectar (Liu *et al*. 2007). In addition, flavonoids (one of the most frequently highlighted chemical classes in our study) influence flower development and pollen fertility (Taylor & Grotewold 2005), while cinnamic acids and derivatives could serve antioxidant properties or indicate their links with volatile biosynthesis (Kikuzaki *et al*. 2002; Maoz *et al*. 2022). Overall, our evolutionary experimental approach provided a holistic understanding of the phytochemical traits selected to cope with limited pollinator access, including chemicals involved in (i) floral and leaf traits (*e.g.* terpenoids, flavonoids), (ii) pollinator attraction (*e.g.* nectar composition), and (iii) plant defence (*e.g.* glucosinolates, in line with selfing promotion).

## 5. Concluding remarks

Despite the central role of phytochemical traits in the structure of plant-pollinator networks, our understanding is still limited to specific chemical classes in flowers and does not consider chemical diversity or the evolutionary dynamics of these traits. Here, we combined an experimental evolution experiment with predictive metabolomics to fill this knowledge gap. Our findings have multiple consequences from chemical ecology to ecosystem dynamics. First, results support the evolutionary capacity of chemodiversity and provide relevant insights into important chemical classes for plant-insect interactions in both flowers and leaves. Second, our results reinforce the value of predictive metabolomics as an ideal tool for studying ecological processes (Díaz *et al*. 2024; Dussarrat *et al*. 2022). Finally, previous studies highlighted that higher biological and chemical diversity of flowering plants enhances pollinator richness and the resilience of plant-pollinator networks (Fornoff *et al*. 2017; Fründ *et al*. 2010). Our study highlights a decrease in chemical richness and diversity of plants when pollinator access is reduced, becoming more specialised to specific pollinators, which could in turn accelerate pollinator decline, pinpointing a consequent risk to ecosystem stability. Changes in plant metabolites due to limited pollinator access may also have consequences on various other interacting organisms, such as herbivores, florivores or pathogens (Hanusch *et al*. 2025).

## Author contributions

EA, LF, CM, and TD conceived the project and designed the experiment. EA, LF and TD realised the sampling. LB, JF and TD performed chemical extractions and GC-MS and LC-MS measurements. RJ and TD pre-processed the data. EA and TD analysed the data. EA, LF, RJ, CM and TD interpreted the results. EA, LF and TD wrote the first draft, and all authors reviewed and edited the manuscript.

## Acknowledgements.

EA and LF thank the University of Zurich and the University Research Priority Program ‘Evolution in action’ for their support. CM and TD acknowledge the financial support from the German Research Foundation (DFG, project MU 1829/28-2). TD acknowledges the Faculty of Biology of Bielefeld University for financial support (Project I-3210-0213-0014, Forschung und wissenschaftlicher Nachwuchs). We also acknowledge the Open Access Publication Fund of Bielefeld University.

## Conflicts of Interest

The authors declare no conflicts of interest.

## Data Availability Statement

Data and metadata are included in supplemental tables and were deposited on MassIVE (MSV000098901, https://doi.org/doi:10.25345/C5BG2HP7R), which includes raw data (GC-MS and LC-MS), non-normalised and normalised GC-MS and LC-MS datasets and R scripts used for this study. Data will be made available upon acceptance of this study.

## Notes

### Competing Interest Statement

The authors have declared no competing interest.

## References

1. Aguirre, L.A. & Junker, R.R. (2024). Floral and pollinator functional diversity mediate network structure along an elevational gradient. Alpine Botany, 134, 193–206.

2. Aizen, M.A., Garibaldi, L.A., Cunningham, S.A. & Klein, A.M. (2009). How much does agriculture depend on pollinators? Lessons from long-term trends in crop production. Annals of Botany, 103, 1579–1588.

3. Artamendi, M., Martin, P.A., Bartomeus, I. & Magrach, A. (2024). Loss of pollinator diversity consistently reduces reproductive success for wild and cultivated plants. Nature Ecology & Evolution, 9, 296–313.

4. Authier, E., Aeschbacher, S. & Frachon, L. (2026). Phenotypic evolutionary response to temporally limited pollinator access in Brassica rapa.

5. Balfour, N.J., Ollerton, J., Castellanos, M.C. & Ratnieks, F.L.W. (2018). British phenological records indicate high diversity and extinction rates among late-summer-flying pollinators. Biological Conservation, 222, 278–283.

6. Bascompte, J. & Jordano, P. (2007). Plant-animal mutualistic networks: the architecture of biodiversity. *Annual Review of Ecology*, Evolution, and Systematics, 38, 567–593.

7. Basnett, S., Krpan, J. & Espíndola, A. (2025). Floral traits and their connection with pollinators and climate. Annals of Botany, 135, 125–140.

8. Biesmeijer, J.C., Roberts, S.P.M., Reemer, M., Ohlemüller, R., Edwards, M., Peeters, T., et al. (2006). Parallel declines in pollinators and insect-pollinated plants in Britain and the Netherlands. Science, 313, 351–354.

9. Borghi, M. & Fernie, A.R. (2017). Floral metabolism of sugars and amino acids: implications for pollinators’ preferences and seed and fruit set. Plant Physiologist, 175, 1510–1524.

10. Bouhaddani, S.E., Uh, H.-W., Jongbloed, G., Hayward, C., Klarić, L., Kiełbasa, S.M., et al. (2018). Integrating omics datasets with the OmicsPLS package. BMC Bioinformatics, 19, 371.

11. Brückner, A. & Heethoff, M. (2017). A chemo-ecologists’ practical guide to compositional data analysis. Chemoecology, 27, 33–46.

12. Burkle, L.A. & Runyon, J.B. (2019). Floral volatiles structure plant–pollinator interactions in a diverse community across the growing season. Functional Ecology, 33, 2116–2129.

13. Calf, O.W., Huber, H., Peters, J.L., Weinhold, A., Poeschl, Y. & van Dam, N.M. (2019). Gastropods and insects prefer different *Solanum dulcamara* chemotypes. Journal of Chemical Ecology, 45, 146–161.

14. Defossez, E., Pitteloud, C., Descombes, P., Glauser, G., Allard, P.-M., Walker, T.W.N., et al. (2021). Spatial and evolutionary predictability of phytochemical diversity. Proceedings of the National Academy of Sciences USA, 118, e2013344118.

15. Deslous, P., Bournonville, C., Decros, G., Okabe, Y., Mauxion, J.-P., Jorly, J., et al. (2021). Overproduction of ascorbic acid impairs pollen fertility in tomato. Journal of Experimental Botany, 72, 3091–3107.

16. Díaz, F.P., Dussarrat, T., Carrasco-Puga, G., Colombié, S., Prigent, S., Decros, G., et al. (2024). Ecological and metabolic implications of the nurse effect of *Maihueniopsis camachoi* in the Atacama Desert. New Phytologist, 241, 1074–1087.

17. Dorey, T. & Schiestl, F.P. (2024). Bee-pollination promotes rapid divergent evolution in plants growing in different soils. Nature Communications, 15, 2703.

18. Dudareva, N., Negre, F., Nagegowda, D.A. & Orlova, I. (2006). Plant volatiles: recent advances and future perspectives. Critical Reviews in Plant Sciences, 25, 417–440.

19. Dussarrat, T., Nilo-Poyanco, R., Moyano, T.C., Prigent, S., Jeffers, T.L., Díaz, F.P., et al. (2024). Phylogenetically diverse wild plant species use common biochemical strategies to thrive in the Atacama Desert. Journal of Experimental Botany, 75, 3596–3611.

20. Dussarrat, T., Prigent, S., Latorre, C., Bernillon, S., Flandin, A., Díaz, F.P., et al. (2022). Predictive metabolomics of multiple Atacama plant species unveils a core set of generic metabolites for extreme climate resilience. New Phytologist, 234, 1614–1628.

21. Dussarrat, T., Schweiger, R., Ziaja, D., Nguyen, T.T.N., Krause, L., Jakobs, R., et al. (2023). Influences of chemotype and parental genotype on metabolic fingerprints of tansy plants uncovered by predictive metabolomics. Scientific Reports, 13, 11645.

22. Eckert, C.G., Kalisz, S., Geber, M.A., Sargent, R., Elle, E., Cheptou, P.-O., et al. (2010). Plant mating systems in a changing world. Trends in Ecology & Evolution, 25, 35–43.

23. Fornoff, F., Klein, A., Hartig, F., Benadi, G., Venjakob, C., Schaefer, H.M., et al. (2017). Functional flower traits and their diversity drive pollinator visitation. Oikos, 126, 1020–1030.

24. Friedman, J., Hastie, T. & Tibshirani, R. (2010). Regularization paths for generalized linear models via coordinate descent. Journal of Statistical Software, 33, 1–22.

25. Fründ, J., Linsenmair, K.E. & Blüthgen, N. (2010). Pollinator diversity and specialization in relation to flower diversity. Oikos, 119, 1581–1590.

26. Gervasi, D.D.L. & Schiestl, F.P. (2017). Real-time divergent evolution in plants driven by pollinators. Nature Communications, 8, 14691.

27. Gu, Z., Gu, L., Eils, R., Schlesner, M. & Brors, B. (2014). circlize implements and enhances circular visualization in R. Bioinformatics, 30, 2811–2812.

28. Hallmann, C.A., Sorg, M., Jongejans, E., Siepel, H., Hofland, N., Schwan, H., et al. (2017). More than 75 percent decline over 27 years in total flying insect biomass in protected areas. PLoS ONE, 12, e0185809.

29. Hanusch, M., Dötterl, S., Larue-Kontić, A.C., Keller, A. & Junker, R.R. (2025). Floral scent chemodiversity is associated with high floral visitor but low bacterial richness on flowers. New Phytologist, 248, 3270–3279.

30. Hegland, S.J., Nielsen, A., Lázaro, A., Bjerknes, A. & Totland, Ø. (2009). How does climate warming affect plant-pollinator interactions? Ecology Letters, 12, 184–195.

31. Hetherington-Rauth, M.C. & Ramírez, S.R. (2016). Evolution and diversity of floral scent chemistry in the euglossine bee-pollinated orchid genus *Gongora*. Annals of Botany, 118, 135–148.

32. Hoballah, M.E., Gübitz, T., Stuurman, J., Broger, L., Barone, M., Mandel, T., et al. (2007). Single gene–mediated shift in pollinator attraction in *Petunia*. The Plant Cell, 19, 779–790.

33. Hoffmann, M.A., Nothias, L.-F., Ludwig, M., Fleischauer, M., Gentry, E.C., Witting, M., et al. (2022). High-confidence structural annotation of metabolites absent from spectral libraries. Nature Biotechnology, 40, 411–421.

34. Hopkins, R.J., Van Dam, N.M. & Van Loon, J.J.A. (2009). Role of glucosinolates in insect-plant relationships and multitrophic interactions. Annual Review of Entomology., 54, 57–83.

35. Imtiaz, H., Arif, Y., Alam, P. & Hayat, S. (2023). Apocarotenoids biosynthesis, signaling regulation, crosstalk with phytohormone, and its role in stress tolerance. Environmental and Experimental Botany, 210, 105337.

36. Jr, F.E.H. (2023). Hmisc: Harrell Miscellaneous.

37. Kikuzaki, H., Hisamoto, M., Hirose, K., Akiyama, K. & Taniguchi, H. (2002). Antioxidant properties of ferulic acid and its related compounds. Journal of Agricultural and Food Chemistry, 50, 2161–2168.

38. Kim, H.W., Wang, M., Leber, C.A., Nothias, L.-F., Reher, R., Kang, K.B., et al. (2021). NPClassifier: a deep neural network-based structural classification tool for natural products. Journal of Natural Products, 84, 2795–2807.

39. Klein, A.-M., Vaissière, B.E., Cane, J.H., Steffan-Dewenter, I., Cunningham, S.A., Kremen, C., et al. (2007). Importance of pollinators in changing landscapes for world crops. Proceedings of the Royal Society B., 274, 303–313.

40. Kleine, S. & Müller, C. (2013). Differences in shoot and root terpenoid profiles and plant responses to fertilisation in Tanacetum vulgare. Phytochemistry, 96, 123–131.

41. Kolde, R. (2019). pheatmap: Pretty Heatmaps.

42. Kopka, J., Schauer, N., Krueger, S., Birkemeyer, C., Usadel, B., Bergmuller, E., et al. (2005).: the Golm Metabolome Database. Bioinformatics, 21, 1635–1638.

43. Larue, A.C., Raguso, R.A. & Junker, R.R. (2016). Experimental manipulation of floral scent bouquets restructures flower–visitor interactions in the field. Journal of Animal Ecology, 85, 396–408.

44. Lemmon, E.W., Huber, M.L., McLinden, M.O., & others. (2010). NIST standard reference database 23. Reference fluid thermodynamic and transport properties (REFPROP), version, 9.

45. Liu, F., Chen, J., Chai, J., Zhang, X., Bai, X., He, D., et al. (2007). Adaptive functions of defensive plant phenolics and a non-linear bee response to nectar components. Functional Ecology, 21, 96–100.

46. Maoz, I., Lewinsohn, E. & Gonda, I. (2022). Amino acids metabolism as a source for aroma volatiles biosynthesis. Current Opinion in Plant Biology, 67, 102221.

47. McCall, A.C., Case, S., Espy, K., Adams, G. & Murphy, S.J. (2018). Leaf herbivory induces resistance against florivores in *Raphanus sativus*. Botany, 96, 337–343.

48. Oksanen, J., Simpson, G.L., Blanchet, F.G., Kindt, R., Legendre, P., Minchin, P.R., et al. (2022). vegan: Community Ecology Package.

49. Pang, Z., Chong, J., Zhou, G., de Lima Morais, D.A., Chang, L., Barrette, M., et al. (2021). MetaboAnalyst 5.0: narrowing the gap between raw spectra and functional insights. Nucleic Acids Research, 49, W388–W396.

50. Parachnowitsch, A. & Junker, R.R. (2021). Working towards a holistic view on flower traits— how floral scents mediate plant-animal interactions in concert with other floral characters. Journal of the Indian Institute of Science, 95:1.

51. Parachnowitsch, A.L. & Manson, J.S. (2015). The chemical ecology of plant-pollinator interactions: recent advances and future directions. Current Opinion in Insect Science, 8, 41–46.

52. Petanidou, T., Kallimanis, A.S., Sgardelis, S.P., Mazaris, A.D., Pantis, J.D. & Waser, N.M. (2014). Variable flowering phenology and pollinator use in a community suggest future phenological mismatch. Acta Oecologica, 59, 104–111.

53. Petrén, H., Anaia, R.A., Aragam, K.S., Bräutigam, A., Eckert, S., Heinen, R., et al. (2024). Understanding the chemodiversity of plants: Quantification, variation and ecological function. Ecological Monographs, 94, e1635.

54. Petrén, H., Köllner, T.G. & Junker, R.R. (2023). Quantifying chemodiversity considering biochemical and structural properties of compounds with the R package CHEMODIV. New Phytologist, 237, 2478–2492.

55. Potts, S.G., Biesmeijer, J.C., Kremen, C., Neumann, P., Schweiger, O. & Kunin, W.E. (2010). Global pollinator declines: trends, impacts and drivers. Trends in Ecology & Evolution, 25, 345–353.

56. R Core Team. (2022). R: A Language and Environment for Statistical Computing. R Foundation for Statistical Computing, Vienna, Austria.

57. Ramos, S.E. & Schiestl, F.P. (2019). Rapid plant evolution driven by the interaction of pollination and herbivory. Science, 364, 193–196.

58. Rusman, Q., Hooiveld-Knoppers, S., Dijksterhuis, M., Bloem, J., Reichelt, M., Dicke, M., et al. (2022). Flowers prepare themselves: leaf and root herbivores induce specific changes in floral phytochemistry with consequences for plant interactions with florivores. New Phytologist, 233, 2548–2560.

59. Sasidharan, R., Grond, S.G., Champion, S., Eilers, E.J. & Müller, C. (2024). Intraspecific plant chemodiversity at the individual and plot levels influences flower visitor groups with consequences for germination success. Functional Ecology, 38, 2665–2678.

60. Sasidharan, R., Junker, R.R., Eilers, E.J. & Müller, C. (2023). Floral volatiles evoke partially similar responses in both florivores and pollinators and are correlated with non-volatile reward chemicals. Annals of Botany, 132, 1–14.

61. Schauer, N., Steinhauser, D., Strelkov, S., Schomburg, D., Allison, G., Moritz, T., et al. (2005). GC–MS libraries for the rapid identification of metabolites in complex biological samples. FEBS Letters, 579, 1332–1337.

62. Schiestl, F.P. (2010). The evolution of floral scent and insect chemical communication. Ecology Letters, 13, 643–656.

63. Schweiger, R., Maidel, A., Renziehausen, T., Schmidt-Schippers, R. & Müller, C. (2023). Effects of drought, subsequent waterlogging and redrying on growth, physiology and metabolism of wheat. Physiologia Plantarum, 175, e13874.

64. Stevenson, P.C., Nicolson, S.W. & Wright, G.A. (2017). Plant secondary metabolites in nectar: impacts on pollinators and ecological functions. Functional Ecology, 31, 65–75.

65. Tay, J.K., Narasimhan, B. & Hastie, T. (2023). Elastic net regularization paths for all generalized linear models. Journal of Statistical Software, 106.

66. Taylor, L.P. & Grotewold, E. (2005). Flavonoids as developmental regulators. Current Opinion in Plant Biology, 8, 317–323.

67. Tenhumberg, B., Dellinger, A.S. & Smith, S.D. (2023). Modelling pollinator and nonpollinator selection on flower colour variation. Journal of Ecology, 111, 746–760.

68. Thomann, M., Imbert, E., Devaux, C. & Cheptou, P.-O. (2013). Flowering plants under global pollinator decline. Trends in Plant Science, 18, 353–359.

69. Thon, F.M., Müller, C. & Wittmann, M.J. (2024). The evolution of chemodiversity in plants—from verbal to quantitative models. Ecology Letters, 27, e14365.

70. Truffault, V., Fry, S.C., Stevens, R.G. & Gautier, H. (2017). Ascorbate degradation in tomato leads to accumulation of oxalate, threonate and oxalyl threonate. The Plant Journal, 89, 996–1008.

71. Wendell, D.L. & Pickard, D. (2007). Teaching human genetics with mustard: rapid cycling *Brassica rapa* (Fast Plants Type) as a model for human genetics in the classroom laboratory. LSE, 6, 179–185.

72. Whitehead, S.R., Bass, E., Corrigan, A., Kessler, A. & Poveda, K. (2021). Interaction diversity explains the maintenance of phytochemical diversity. Ecology Letters, 24, 1205–1214.

73. Wickham, H. (2016). ggplot2: Elegant graphics for data analysis. Springer-Verlag New York.

74. Ziaja, D. & Müller, C. (2023). Intraspecific chemodiversity provides plant individual– and neighbourhood-mediated associational resistance towards aphids. Frontiers in Plant Science, 14, 1145918.

75. Zu, P. & Schiestl, F.P. (2017). The effects of becoming taller: direct and pleiotropic effects of artificial selection on plant height in *Brassica rapa*. The Plant Journal, 89, 1009–1019.

