## Supplemental Figures for "Metabolism and chemical diversity evolve in response to pollinator availability"

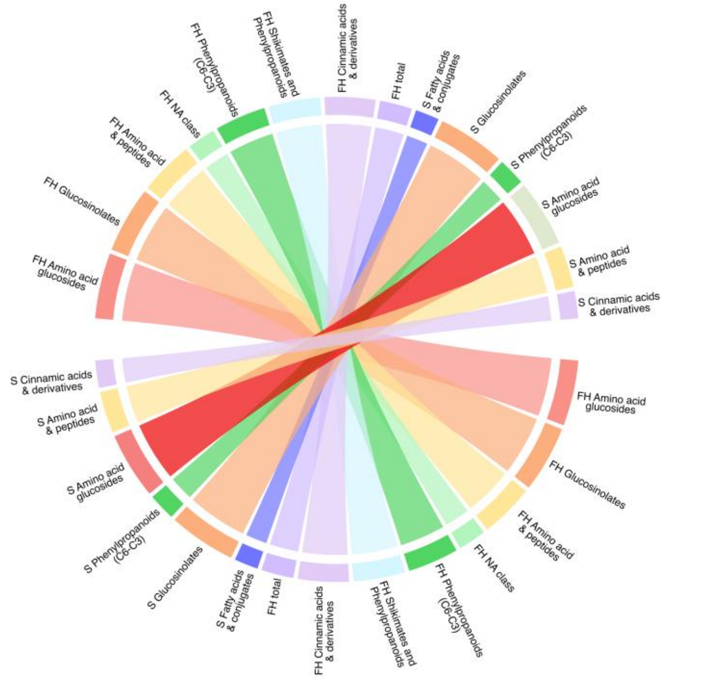


**Fig. S1 | Correlation between chemical indices of flowers and leaves.** Circos plot representing the significant correlations ($P < 0.05$, FDR, Pearson correlation) between chemical indices of flowers (upper part of the figure) and leaves (lower part of the figure). Links are proportional to the correlation (R) and range from 0.27 to 0.77 (all significant correlations were positive). Correlations between chemical classes (*e.g.* correlation between "Amino acid glucoside" and "glucosinolates") were excluded from the figure. *S*: Shannon diversity ; *FH*: Functional Hill diversity.

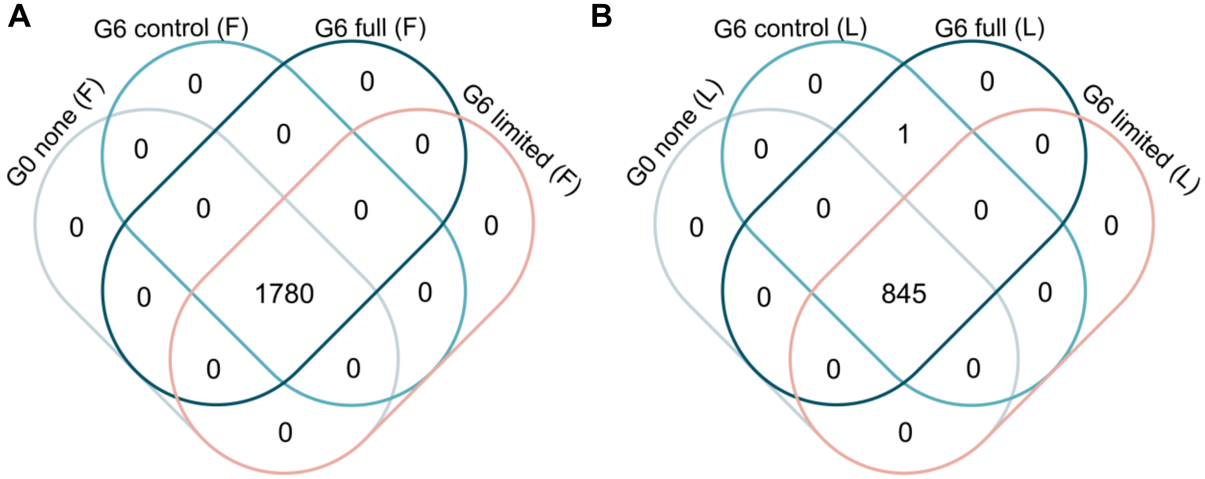


**Fig. S2 | Stability of detected ions between classes in flowers and leaves. A-B.** Venn diagram of detected ions after pre-processing steps in flowers (A) or leaves (B).

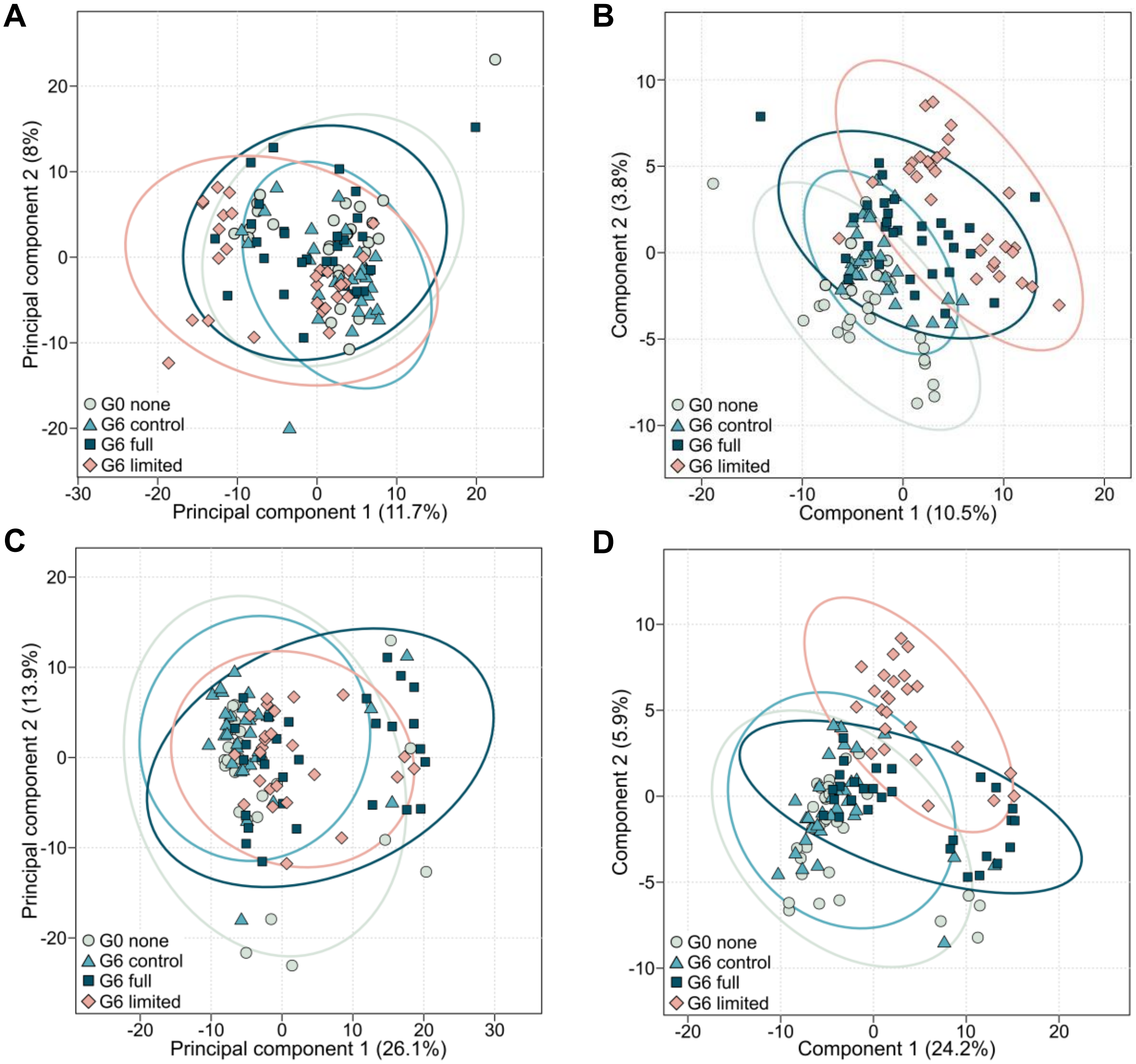


**Fig. S3 | Discriminant capacity of flower and leaf metabolism on dataset 1. A-B.** Principal component analysis (PCA) (A) and partial least squares discriminant analysis (PLS-DA) (B) using pre-processed features from flower. **C-D**. PCA (C) and PLS-DA (D) using pre-processed features from leaves.

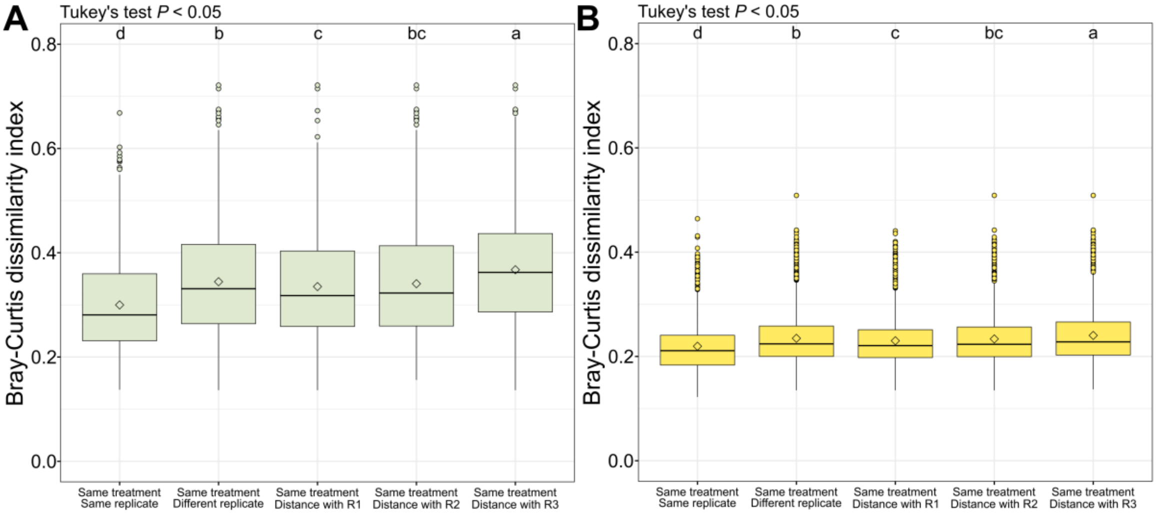


**Fig. S4 | Bray-Curtis dissimilarity between replicates.** Variation of Bray-Curtis dissimilarity between replicates 1, 2 and 3 using LC-MS data in leaves (A) and flowers (B).

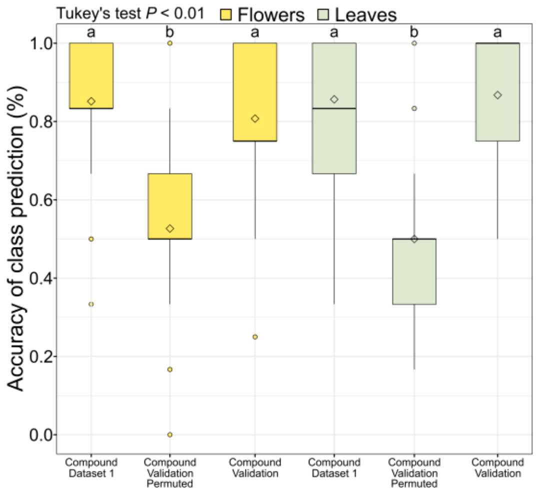


**Fig. S5 | Capacity to predict the type of pollination using leaf or flower metabolism.** *$R^2$* of 100 models to test the capacity of flower and leaf metabolism to distinguish between the classes G6_control (*G6_c*, where pollination was performed by hand) and G6_full (*G6_f*, where pollination was performed by natural pollinators). Only the best 5% predictors of these classes were used to perform the model (*i.e.* 89 features). For validation set, model equation was developed on dataset 1 and directly applied to dataset 2. Significant variation in predictive score between modelling condition were accessed using Tukey's test ($P < 0.01$).

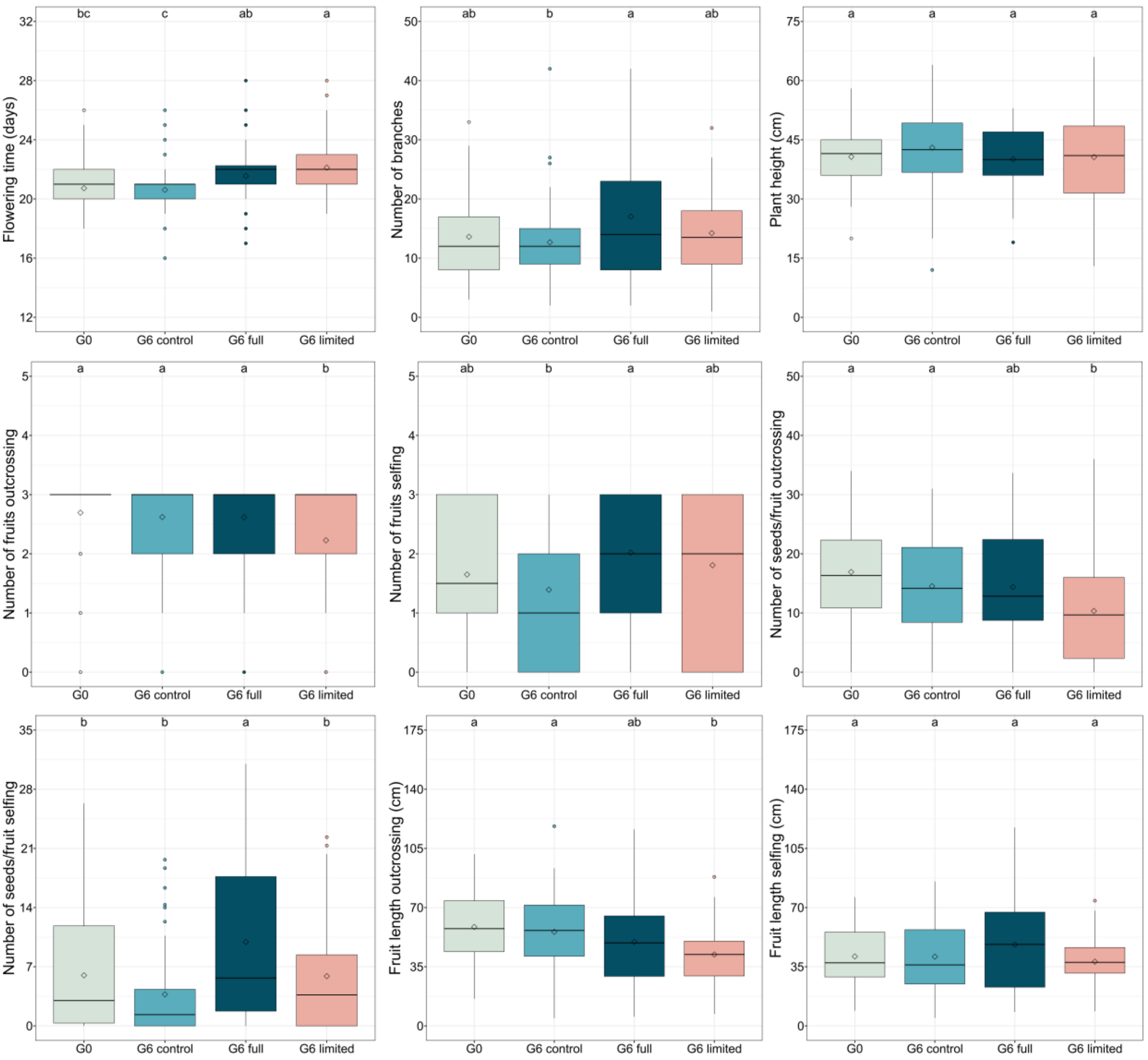


**Fig. S6 | Variation in phenotypic traits in response to pollinator availability.** Significant differences can be seen by the letters in the boxplots representing the results of the Tukey's tests, $P < 0.05$.
